# Autoaggregation in *Streptococcus intermedius* is driven by the Pel polysaccharide

**DOI:** 10.1101/2024.04.20.589940

**Authors:** Deepa Raju, Siobhán A. Turner, Karla Castro, Gregory B. Whitfield, Daphnée LaMarche, Sahil Mahajan, Roland Pfoh, François Le Mauff, Maju Joe, Susmita Sarkar, Todd L. Lowary, Donald C Sheppard, Daniel J. Wozniak, Michael G. Surette, P. Lynne Howell

## Abstract

The Streptococcus Milleri Group (SMG) comprising of *Streptococcus intermedius, anginosus* and *constellatus* are commensal bacteria commonly found in healthy individuals. These bacteria are increasingly being recognized as opportunistic pathogens that can cause purulent infections at sterile body sites and have also been identified in the sputum of individuals with cystic fibrosis. Although the mechanisms of conversion to opportunistic pathogens are not well understood, auto-aggregation is a key driver of biofilm adhesion and cohesion in many Streptococci and Staphylococci. Here, we identify a gene cluster in the *S. intermedius* genome with significant homology to the *pel* operons in *Bacillus cereus* and *Pseudomonas aeruginosa*, which are required for Pel exopolysaccharide production and biofilm formation in these species. Characterization of a panel of clinical *S. intermedius* strains identified a range of aggregating phenotypes. Analysis of the *pel* operon in the hyper-aggregating C1365 strain revealed that each of the canonical *pelDEA_DA_FG* genes, but not the four additional genes are required for aggregation. Further, we demonstrate that C1365 produces a GalNAc-rich exopolysaccharide and that aggregates can be disrupted by the α1,4 *N-*acetylgalactosaminidases, PelA and Sph3, but not other glycoside hydrolases, proteinase K or DNase I. Using an abscess model of mouse infection, we show that Pel driven aggregation leads to longer lasting infections, and that lack of Pel allows for the bacteria to be cleared more effectively. The polymer also affects how the bacteria interacts with the host immune system. Collectively, our data suggest that the *pel* operon has relevancy to *S. intermedius* pathogenicity.

## INTRODUCTION

Bacteria most commonly reside in multicellular communities called biofilms. Biofilms can form on medical devices, such as catheters, feeding tubes, and ventilators as well as on mucosal surfaces within the body and are predicted to be involved in ∼80% of all chronic microbial infections (1–3). Biofilm-embedded bacteria are typically more tolerant to immunological, chemical, and mechanical insults compared to their planktonic counterparts (1, 4, 5). This makes the treatment of biofilm-associated infections extremely challenging and the capacity to form a biofilm is a major virulence factor.

Biofilms can be surface attached or exist as multicellular aggregates in suspension (6–8). There is increasing evidence that suggests aggregates formed in media or host fluids exhibit the same characteristics as surface attached biofilms and are increasingly being recognized for their role in pathogenicity (6, 9–12). Multiple chronic infections have been linked to aggregates including chronic infections in cystic fibrosis (7, 13), dermal wounds (9, 11) and otitis media (14). Bacteria in both surface attached biofilms and aggregates are recalcitrant to antibiotic treatment and are protected from the immune system. These advantages are enabled, at least in part, by the presence of an extracellular matrix made of various components including water, lipids, proteins, extracellular DNA (eDNA), RNA, membrane vesicles, phages, and exopolysaccharides.

One important matrix component in both Gram-negative and Gram-positive bacteria is the Pel polysaccharide (15–17). This polysaccharide has been well-studied in *P. aeruginosa* (18) where it is required for pellicle formation at the air liquid interface and the bacterium’s characteristic wrinkly colony morphology (19, 20). Pel is also critical for cell-to-cell adhesion and formation of aggregates (21). Within the extracellular matrix, Pel exists in two forms: a cell associated and cell free form. A recent study suggests that the two forms of the polymer play distinct roles within the biofilm, with the cell free form contributing the biomechanical properties of the biofilm and decreased virulence of *P. aeruginosa* in *Drosophila melanogaster* and *Caenorhabditis elegans* infection models (19). While the structure of the cell associated form has yet to be determined, the cell free form is a homopolymer of partially de-*N-*acetylated α1, 4 linked *N*-acetylgalactosamine (GalNAc), which consists predominantly of a dimeric repeat of GalNAc and galactosamine (GalN) (22). Deacetylation of the polymer renders Pel cationic and this modification is required for biofilm formation (21). Pel’s cationic charge facilitates its interaction with eDNA(21) and host derived anionic polymers in CF sputum (23); interactions which help stabilize the structural core of the biofilm (18, 23). Pel also interacts with protein components, such as CdrA, and enhances biofilm aggregation (24, 25). The presence of Pel in the extracellular matrix has been shown to increase bacterial tolerance to antibiotic treatment; an effect that is further enhanced by its binding to eDNA (23, 26).

All of the genes in the *P. aeruginosa pelABCDEFG* operon are essential for biofilm and aggregate formation (27–31). PelDEFG form an inner membrane complex that enables polymerization and export of the Pel polymer across the cytoplasmic membrane (32). This biosynthetic complex is activated by the binding of cyclic di-guanosine monophosphate (c-di-GMP) to PelD (33, 34), where upon the glycosyltransferase PelF uses UDP-*N*-acetylgalactosamine as a substrate for Pel synthesis (28). PelE is a predicted protein interaction module, while PelG is the predicted translocase. PelB and PelC help guide the Pel polymer across the periplasm and are required for its export into the extracellular space (30, 35). PelA is a periplasmic modification enzyme that exhibits both glycoside hydrolase and deacetylase activities (19, 29, 36). These activities are required for the production of cell free Pel and Pel deacetylation, respectively (19, 22, 29).

Using a computational pipeline that allows for an unbiased identification of functionally related gene clusters, we recently discovered a *pel-*like gene cluster, *pelDEA_DA_FG,* in a wide variety of Gram-positive bacterial species, including Streptococcal species, and have demonstrated that the GalNAc rich Pel-like polymer produced by this gene cluster is required for biofilm formation in *Bacillus cereus* ATCC 10987 (15, 16). The *pel*-like gene cluster found in Gram-positive bacteria contains the core *Pseudomonas* homologs, *pelEFG,* and a truncated *pelA* homologue, referred to herein is as *pelA_DA_*, which lacks the hydrolase domain found in the *P. aeruginosa* counterpart. While not part of the *pel* gene cluster, a glycoside hydrolase is frequently found divergently encoded up-stream or down-stream in Gram-positive species. In *B. cereus*, similar to *P. aeruginosa,* deletion of the up-stream glycoside hydrolase resulted in increased biofilm biomass (16). The *pelD* gene exhibits the largest divergence across species.

In Streptococcus genomes, including *S. intermedius*, the *pel*-like gene cluster diverges from the canonical Gram-negative *P. aeruginosa pelABCDEFG* and Gram-positive *B. cereus pelDEA_DA_FG* operons in a number of ways including the presence of four additional genes whose function in Pel biosynthesis and biofilm formation are currently unknown (Fig. 1). The most prominent difference is in *pelD* (31, 33). *P. aeruginosa* and *B. cereus pelD* encode proteins, *Pa*PelD and *Bc*PelD, with 4 transmembrane (TM) domains, and cytoplasmic GAF and GGDEF domains, which bind c-di-GMP to regulate Pel biosynthesis (33, 34). The TM-GAF-GGDEF domain architecture observed in *Pa*PelD and *Bc*PelD is conserved in other *Bacillus* species as well as *Halobacillus, Pontibacillus, Salimicrobium, Marinococcus, Exiguobacterium* spp (16, 17). In contrast, in *Streptococcus* spp the *pelD* homologue encodes a highly divergent protein (33). While *Si*PelD has the TM and GAF domains, it lacks the GGDEF domain suggesting that the Pel protein complex is not regulated by c-di-GMP in these species. In addition, *Si*PelD contains an additional N-terminal domain that is predicted to be a degenerate short chain dehydrogenase/reductase (SDR) (17). In some Gram-positive species including some *Clostridial*, *Paenibacillus, Neobactillus* and *Neobacillus* spp, PelD contains both the GGDEF domain as well as a functional N-terminal SDR domain, while others, including *Bifidobacterium* contain the GGDEF domain and a degenerate SDR domain. The active SDR domain is predicted to be an epimerase, potentially involved in production of the nucleotide-sugar precursor. The function of the degenerate SDR domain is currently unknown.

**Fig. 1.**
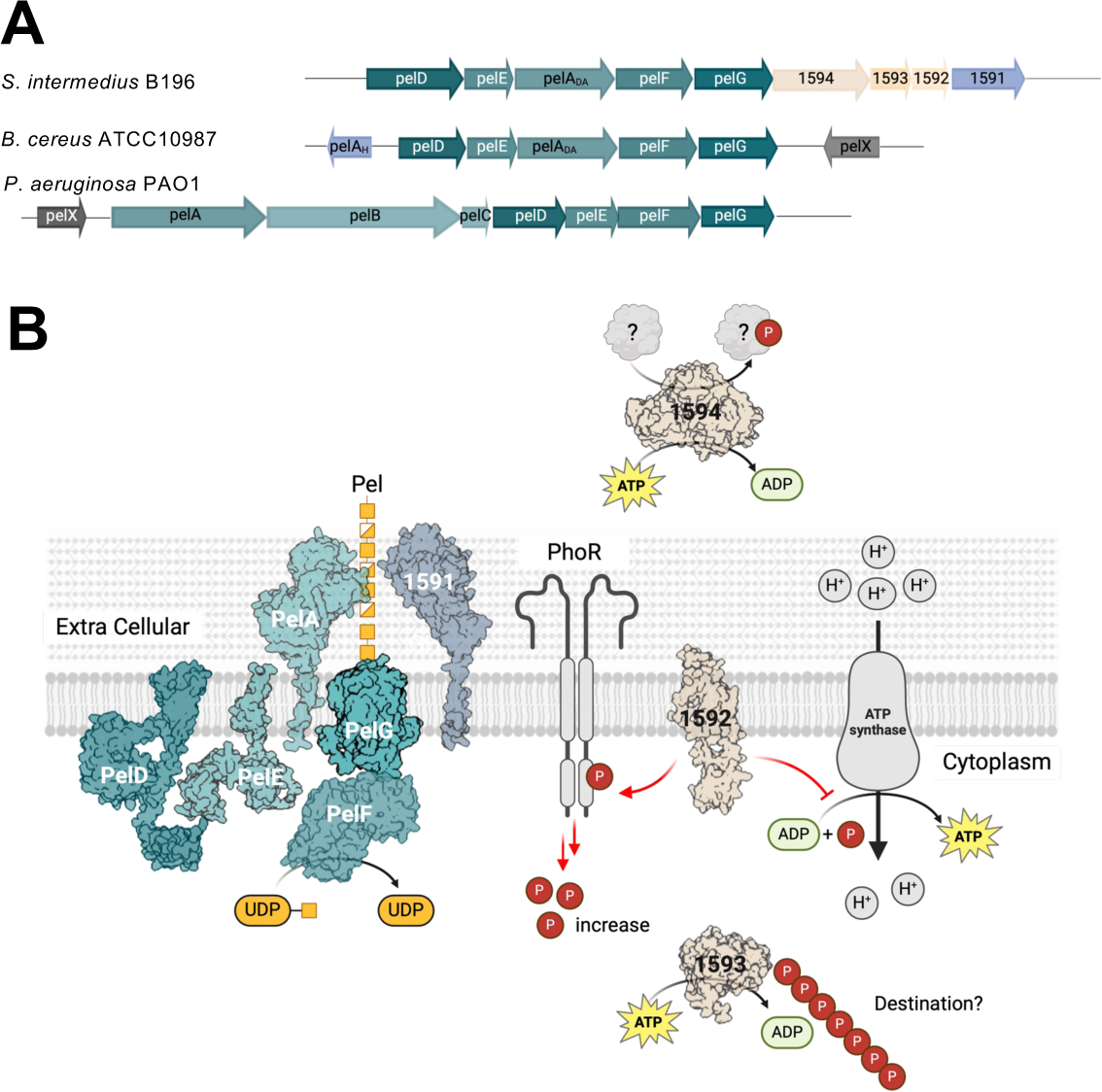
Model of the Pel biosynthetic complex in *S. intermedius*. (A) Schematic representation of the *S. intermedius, B. cereus* and *P. aeruginosa pel* operons. Additional genes in the *pel* operon are indicated in wheat or pale blue. (B) Proposed model for the *S. intermedius* pel biosynthesis complex based on its homology with *P. aeruginosa.* The surface representation of the AF2 model of each protein and potential roles of the additional genes in the *S. intermediate pel* operon are depicted.

This partially divergent Pel gene cluster is found in the larger Viridans group of Streptococci. The Streptococcus milleri group (SMG) is a part of this larger group and consists of *Streptococcus intermedius*, *Streptococcus constellatus* and *Streptococcus anginosus*. These species are normally considered commensal bacteria that commonly reside in the oral, gastrointestinal, and urogenital tracts of healthy individuals (37) but with the advent of better detection and diagnostic techniques, their relevance as opportunistic pathogens is increasingly being recognized (38, 39). These bacteria cause purulent infections at sterile body sites (40, 41) and were found to be the most common cause of invasive pyogenic streptococcal infections in the Calgary Health Region between 1999 and 2004 (42). More recently, these species have been identified as significant pathogens in the lungs of individuals with cystic fibrosis (43–49). Despite their role as opportunistic human pathogens, little is known about how SMG transition from harmless commensals to pathogens, and which environmental signals induce this change in lifestyle.

Given the role of Pel in other species and lack of any other identifiable exopolysaccharide biosynthetic operons we hypothesized that Pel may play a role in biofilm formation in SMG species and thus contribute to their pathogenicity and virulence. Using *S. intermedius* as our model organism, we established a role for Pel in aggregate formation in these species. Screening a panel of clinical *S. intermedius* isolates, we discovered a range of aggregation phenotypes across the eleven strains tested. We found that all the canonical *pel* genes were essential for aggregation and the presence of the polymer was associated with persistence of infection in a mouse abscess model. Additionally, we demonstrate that under the conditions tested, the four additional genes present in the operon do not directly affect polymer production or aggregate formation.

## MATERIALS AND METHODS

### Strains and media

*S. intermedius* strains were cultured at 37 °C, 5% CO_2_ using THY broth (37 g/L Todd Hewitt Broth, 0.5% (w/v) yeast extract), or THY agar (37 g/L Todd Hewitt Broth, 0.5% (w/v) yeast extract, 1.5% agar). Antibiotic selection was performed by adding 250 µg/mL kanamycin as required. A list of the clinical strains used in this study and the source they were isolated from is presented in Table 1.

**Table 1:** List of *S. intermedius* strains used in this study.

| **Strain designation** | **Anatomical Source** | **Anatomical Source** |
| --- | --- | --- |
| C1365 | Blood | Calgary, AB, Canada |
| C1369S | Blood | Calgary, AB, Canada |
| C1369R | Blood | Calgary, AB, Canada |
| C1377 | Blood | Calgary, AB, Canada |
| C1390 | Brain | Calgary, AB, Canada |
| C260 | Invasive | Calgary, AB, Canada |
| M60R | Exacerbation | Calgary, AB, Canada |
| B196 | Hip abscess | Calgary, AB, Canada |
| SAG05 | Neck lymph node | Toronto, ON, Canada |
| AP1 | Pleural empyema | Arizona, USA |
| AP2 | Pleural empyema | Arizona, USA |

### Gene cluster analysis

The online server CAGECAT (50) was used to evaluate how widespread the additional genes in the *pel* operon, *SIR_1592-1594,* are and whether this set of genes exists in this combination independent from the *pel* operon. NCBI accession gene numbers for the target genes were provided to cblaster and cblaster run using its default parameters. We examined two cases, firstly the co-occurrence of *S. intermedius* genes *SIR_1592-1594* and secondly all genes in the *pel* operon SIR_1591-1599. To avoid duplicate entries of the same species, only NCBI sequences with NZ_ and NC_ prefixes were kept for analysis.

### Generation of mutant strains

Mutants were generated by replacing the target gene with an antibiotic resistance cassette as described previously (51, 52). Primers and plasmids used in the study are listed in Table S1. All primers were ordered from Sigma Aldrich. *S. intermedius* mutants were generated by replacing the gene to be deleted with a cassette (52, 53) conferring resistance to kanamycin as previously described (53). Briefly, the antibiotic resistance cassette was cloned between ∼1000 bp of sequence homologous to the regions flanking the gene to be deleted. The DNA fragment containing the cassette and flanking sequences was then linearized by restriction digest, gel purified, and 50-100 ng of the purified fragment was added to overnight cultures diluted to OD_600_ ∼ 0.08 in 3 mL THY and incubated for 1 h at 37 °C, 5% CO_2_ before adding DNA and 500 ng/mL competence stimulating peptide (52, 53). Cultures were incubated for a further 3-4 h before plating on THY agar containing the relevant antibiotic. The deletion and presence of the antibiotic cassette were further confirmed by PCR. The knockout constructs encoding 1000 bp upstream and downstream with a Kanamycin insertion for *pelF* and *SIR_1591* were designed, codon optimized and were synthesized by BioBasic. All constructs were sequence verified by The Centre for Applied Genomics (TCAG) at The Hospital for Sick Children.

### Analysis of surface adherence

Surface adherence of the 11 clinical strains and the mutants was determined using a crystal violet assay as described previously (54, 55). Overnight cultures were diluted to OD_600_ ∼ 0.005 and incubated for 24 h. Unattached cells were removed by submerging plates twice in water and discarding the run-off. Adherent cells were stained in 0.1% Crystal Violet for 30 min and excess stain removed by submerging plates twice in water. Staining was quantified by eluting Crystal Violet in 30% acetic acid and measuring absorbance at 550 nm.

### Aggregation assays

Aggregation assays were performed after the *S. intermedius* strains were grown overnight at 37 °C. Cultures were centrifuged at 3000 × *g* for 5 min. Media was removed by resuspending cells in 1 mL aggregation buffer (1 mM Tris, 2 mM CaCl_2_, 3 mM MgCl_2_, 150 mM NaCl, buffered to pH 7.4 with HCl) and centrifuging at 3000 × *g* for 5 min. Cells were resuspended in 1 mL aggregation buffer and diluted to OD_600_ ∼ 0.5 - 1.0 in 1 mL aggregation buffer in polyurethane cuvettes. Cuvettes were incubated at room temperature without agitation and the OD_600_ was measured at regular intervals up to 180 mins.

For the aggregate disruption assays using the different enzymes, aggregates were disrupted by adding a concentration range of each of the enzymes to the aggregation buffer and incubating at room temperature with mixing for 1 h. The recombinant hydrolases PslG, PelA, Sph3, Ega3 and Dispersin B were purified using nickel affinity chromatography as described previously (54, 56–59). Chitinase, cellulase and Proteinase K were purchased from Sigma Aldrich, and DNase I was from Biobasic. After incubation with the enzymes for 1h, the cuvettes were further left standing at room temperature and aggregation assayed as described above. The OD 600 over time at different concentrations of the enzymes was analyzed using Prism and the EC50 values calculated as previously described (54).

### Scanning electron microscopy

Overnight cultures were centrifuged at 3000 × *g* for 5 min. Media was removed by resuspending cells in 1 mL PBS and centrifuging at 3000 × *g* for 5 min. Cells were resuspended in 1 mL PBS and diluted to OD_600_ ∼ 1.0. Bacterial samples were prepared for analysis by scanning electron microscopy (SEM) using critical point drying and gold sputtering as previously described (16). Briefly, bacteria were transferred onto coverslips and fixed with 2.5% glutaraldehyde. The coverslips were washed with PBS and dehydrated using a series of ethanol washes at 50, 70, 90 and 100% ethanol for 15 min each. Samples were then critical point dried by replacing the ethanol with CO_2_ and sputter coated in gold. The sample processing was completed by the Nanoscale Biomedical Imaging Facility at The Hospital of Sick Children (Toronto, Ontario) and images were obtained using a Scanning Electron Microscope (FEI 30Kv voltage model XL-30 SEM with a tungsten gun and secondary electron detector) at The Hospital for Sick Children.

### Generation of GalNAc-specific monoclonal antibody

Mice were immunized with a bovine serum albumin (BSA)-(GalNAc)_3_ glycoconjugate followed by natural infection with *Aspergillus fumigatus* Af293 to further mature the immune response. Mice sera were screened for the production of antibodies directed against the oligosaccharide, and immortalized hybridomas were created from their splenocytes. Monoclonal antibody candidates were then screened for reactivity against BSA-(GalNAc)_3_. To identify clones that specifically recognize the GalNAc found in the *P. aeruginosa* Pel and *A. fumigatus* galactosaminogalactan (GAG) exopolysaccharides, Pel and GAG sufficient and deficient culture supernatants of *P. aeruginosa* PA14 and *A. fumigatus* Af293 were used.

### Dot blot analysis

Dot blots using the α-(GalNAc)_3_ antibody were performed as follows. Cells were harvested by centrifugation (4,000 × *g* for 5 min) from 5 mL of *S. intermedius* cultures grown overnight at 37 °C in THY broth. Cell were resuspended in 1 mL aggregation buffer at an optimized OD of 2.0 and centrifuged at 4000 × *g* for 5 min. The supernatant was discarded, and cell pellets were resuspended in 100 μL of 0.5 M EDTA, pH 8.0. Cells were boiled for 20 min with occasional vortexing, and centrifuged (16,000 × *g* for 10 min) to harvest the supernatant containing cell associated Pel. Cell-associated Pel was treated with proteinase K (final concentration, 0.5 mg/mL) for 60 min at 60 °C, followed by 30 min at 80 °C to inactivate proteinase K. For the immunoblots, 5 μL of cell associated Pel, prepared as described above, was pipetted onto a nitrocellulose membrane and left to air dry for 10 min. The membrane was blocked with 5% (w/v) skim milk in Tris-buffered saline with Tween-20 (TBS-T) for 1 h at room temperature and probed with adsorbed α-(GalNAc)_3_ at a 1:1000 dilution in 1% (w/v) skim milk in TBS-T overnight at 4 °C with shaking. Blots were washed three times for 5 min each with TBS-T, probed with goat α-rabbit HRP-conjugated secondary antibody (Bio-Rad) at a 1:2000 dilution in TBS-T for 45 min at room temperature with shaking, and washed again. All immunoblots were developed using SuperSignal West Pico (Thermo Scientific) following the manufacturer’s recommendations.

### *In vivo* murine skin infection model

6- to 8-week-old female BALB/c mice were obtained from Charles River Laboratories and utilized for establishing the skin abscess model. Prior to infection, the mice were anesthetized with isoflurane, and their dorsal backs were shaved. The mice were subsequently intradermally inoculated with *S. intermedius* C1365 wild type or τ1*pelF* strains in 100 ml of PBS. Bacteria for infection were grown overnight in THY broth as described above. On the day of infection, the OD_600_ was determined, and cultures diluted to obtain 10^8^ bacteria/100μl of PBS. Mice were carefully monitored for any signs of skin infection. On day 6, the mice were euthanized, and the abscesses were measured, excised and their weights recorded. Abscesses from 5 mice in each group were homogenized and dilutions plated for enumerating the bacterial burden on THY plates. The plates were incubated overnight at 37 °C in a 5% CO_2_ incubator and the CFU/abscess calculated the next day. All data points were plotted using Graph Pad Prism and an unpaired T test was used for statistical analysis.

### PBMC isolation and cytokine quantification

Peripheral blood mononuclear cells (PBMCs) were recovered from heparinized blood via density gradient centrifugation using Ficoll-Plaque separation media (GE Healthcare) and Leucosep tubes (Greiner Bio-One). PBMCs frozen at −120 °C were quickly thawed, washed, and seeded to a density of 1 × 10^5^ cells/well in 200 μL of RPMI-1640 supplemented with 10 % fetal bovine serum (FBS; Gibco), 100 U/mL penicillin G, 100 μg/mL streptomycin and 2 mM L-glutamine using round-bottom 96 well plates (Corning). PBMCs were activated using heat-killed bacteria at a multiplicity of infection (MOI) of 1:1, 10 ng/mL of LPS (Escherichia coli O55:B5, Sigma-Aldrich) or with supernatants of *S. intermedius* cultures for 24 h at 37 °C in the presence of 5% CO_2_. Cells were subsequently centrifuged at 1,500 × *g* for 5 min and supernatants were recovered and stored at −20 °C until further use. PBMCs viability was confirmed using a CytoTox96 non-radioactive cytotoxicity assay kit (Promega). For cytokine quantification experiments, triplicate reactions were pooled from individual PBMC stimulation experiments and performed in duplicate. Cytokine production was normalized with the LPS-PBMC control to correct for donor and experimental heterogeneity.

## RESULTS

### The *S. intermedius pel* operon contains four additional genes

Our analysis of the *S. intermedius* genome revealed four additional genes as part of its *pel* operon, including a putative glycoside hydrolase, SIR_1591 (Fig. 1). Glycoside hydrolases are a common feature of exopolysaccharide biosynthetic systems (60) and in *P. aeruginosa* the hydrolase activity of PelA has been associated with both production of cell free Pel, as well as matrix disruption and dispersal (19, 61). Our bioinformatics analyses suggest that SIR_1591 will be found on the extracellular surface of the bacterium, in keeping with the identification of a homologue of this protein in the secretome of *Streptococcus gordonii* (62) and the location of the mature Pel polymer. At present it is unclear whether this attachment is *via* a lipid anchor or a transmembrane helix. While SignalP v5 predicted with 70% probability that there is a Sec/SPII lipoprotein signal sequence and cleavage site between residues Ala28 and Cys29, the UniProt server suggests that residues 13-34 form a transmembrane domain. In addition, the AlphaFold2 (AF2) model predicts that SIR_1591 is a two-domain protein with residues 60-350 forming a TIM barrel domain that the DALI and Foldseek (63, 64) servers suggest has structural similarity to beta-amylases and beta-galactosidases. The AF2 model also suggests that residues 44-52 and 351-454 form a seven β-strand domain of unknown function (DUF4832). Homologues of this domain have been found on the C-terminal side of the glycoside hydrolases and have structural similarity to carbohydrate binding domains (65). While the exact role of the SIR_1591 glycoside hydrolase in Pel production remains to be determined, we anticipate that – like *P. aeruginosa* PelA – it could play multiple roles in biofilm formation (19, 29, 35).

Additional bioinformatics analyses reveal that a homologue of the first of the additional three genes, SIR_1594, is also found in the secretome of *Streptococcus gordonii* (62) and has been identified as a putative surface protein in *Streptococcus salivarious* F60-1 (66). Our analyses suggest that this gene encodes a secreted protein whose N-terminal region (residues 62-472) has significant with homology (22% identity) to *Bacillus subtilis* and *Bacillus cereus* CotH, a serine kinase involved in the regulation of spore coat formation. As *Streptococci* do not sporulate and SIR_1594 appears to have the residues required for catalysis, we hypothesize that the kinase function of SIR_1594 has perhaps been re-purposed in this species. The C-terminal region of SIR_1594, residues 474-569, is predicted to contain a β-sandwich fibronectin type-III domain (FN3). FN3 domains are found in a wide variety of proteins including extracellular matrix proteins, cell-surface receptors, and enzymes. While sites of interaction with other molecules have been mapped to the Arg-Gly-Asp (RGD) sequence motif found in various FN3 domains, SIR_1594 lacks this motif, and it is currently unclear what role this domain may play in Pel biosynthesis. We speculated that this domain might assist in targeting a cell-surface or extracellular matrix protein of the host for phosphorylation by the serine kinase-containing domain.

The second additional gene, SIR_1593, encodes a putative cytoplasmic protein, which the AF2 model predicts is structurally similar to the polyphosphate (polyP) polymerase subunit found within the membrane vacuolar transport chaperone (VTC) complex. The VTC complex produces polyP and releases it into intracellular vacuoles (67, 68). The SIR_1593 polymerase is predicted to be comprised of a tunnel-shaped domain formed by antiparallel β-stands surrounded by five α-helices with the walls of the tunnel lined by conserved basic residues. PolyP is an anionic polymer with a diverse array of functions in bacteria including phosphate storage, stress resistance, biofilm formation and virulence (69). Bacterial polyphosphates have also been associated with better survival in human macrophages (70). The lack of the other protein components of the VTC complex in the *pel* operon and the presence of the required catalytic residues in SIR_1593 suggests that the polyP produced may remain in the cytoplasm. What the role of SIR_1593 and polyP are in *S. intermedius* Pel biosynthesis and biofilm formation remains to be determined.

The third additional gene in the *S. intermedius pel* operon, SIR_1952, is predicted to encode a transmembrane protein with 4 TM segments (residues 22-143) and an intracellular C-terminal ACT domain. ACT domains are found in many different proteins involved with metabolism. They are generally thought to be a ligand-binding domain with a role in allosteric regulation of protein function (71, 72). The Conserved Domain Database (CDD) (73) assigns the transmembrane domain of SIR_1952 to the MgtC family. MgtC was originally found in the same operon as the Mg^2+^ transporter MgtB, however it does not transport Mg^2+^, but instead is involved in ATP/H+ regulation in *Salmonella enterica* by inhibiting its own ATP-synthase (74), a process that appears to be related to long-term virulence. In addition, *S. enterica* MgtC also enables intra-macrophage survival by deregulating its phosphate metabolism via targeting PhoR (75). *Salmonella enterica* MgtC contains an N-terminal TM domain and a C-terminal ACT domain and has 17% sequence identity to SIR_1952. The function of this MgtC homologue in *S. intermedius* is currently unknown.

As previous analyses had identified *SIR_1591-1594* downstream of the *pel* operon in a wide variety of *Streptococcal* spp, and some *Bifidobacteria, Clostridia* and *Butyrivibrio* spp (16), we examined whether the combination of these genes is present in other species outside the *pel* operon. A gene cluster analysis using the CAGECAT server (50) was not always able to find all of the expected genes in the canonical *pelDEA_DA_EG* operon. This may be the result of low sequence similarity, low query cover due to the absence of a domain, gaps between genes or actual loss of genes. Focusing our analysis on species for which the entire *pelDEA_DA_EG* operon was found, the analysis revealed that *SIR_1592-1594* also occurred downstream of *pelG* in *Abiotrophia, Blautia, Dorea, Mediterranei* and *Ruminococcus* species (Table S2), while *SIR_1591* appears to occur independently of *SIR_1592-1594* and can be found both downstream and upstream of the *pel* operon. The occurrence of *SIR_1592-1594* was most common in *pel* operons that contain an inactive SDR domain in PelD. Our analysis was unable to find this combination of *SIR_1592-1594* genes outside the *pel* operon (Table S3) suggesting that these genes could play a role in Pel production and/or in infection as SIR_1593 and SIR_1594 have potential roles associated with bacterial survival in macrophages.

### *S. intermedius* clinical strains have variable surface adherent biofilm phenotypes

To determine the role of the *pel* operon in biofilm formation in *S. intermedius* we gathered a collection of eleven clinical isolates from different anatomical sites and geographical locations (Table 1) and tested their ability to adhere to poly-lysine coated polystyrene multi-cell plates using a crystal violet adherence assay (Fig. 2A). We found significant variability in the capacity of the different *S. intermedius* strains to form adherent biofilms under the conditions tested. Some strains, e.g., B196, C1390, SAG05, C1369R, C1369S, C260 and C1377, consistently had high levels of biofilm biomass, while others, such as M60R, AP1, AP2 and C1365 adhered poorly. To determine whether adherence and the levels of biofilm biomass observed were dependent on the Pel polysaccharide, we constructed *pelF* deletion mutants (*ΔpelF*) for each of the clinical isolates and tested these strains using our crystal violet adherence assay. Of the strains that strongly adhered to the plate, deletion of *pelF* in B196, C1369S, C1369R, and SAG05 did not significantly affect their ability to attach to the surface (Fig. S1). Two strongly adhering strains, C1377 and C1390, showed significant differences when *pelF* was deleted but they still attached significantly better than the non-surface adherent M60R, AP1, AP2 and C1365 strains. Deletion of *pelF* in the non-surface adherent strains did not change their non adherent phenotype. These data suggests that while Pel may play a partial role in surface adherence, other components such as eDNA or proteins influence the formation of adherent biofilms more than Pel. In contrast to Pel’s established role in *P. aeruginosa* and *B. cereus* (16, 18), these data suggests that Pel biosynthesis does not drive surface adherence in *S. intermedius*.

**Fig. 2.**
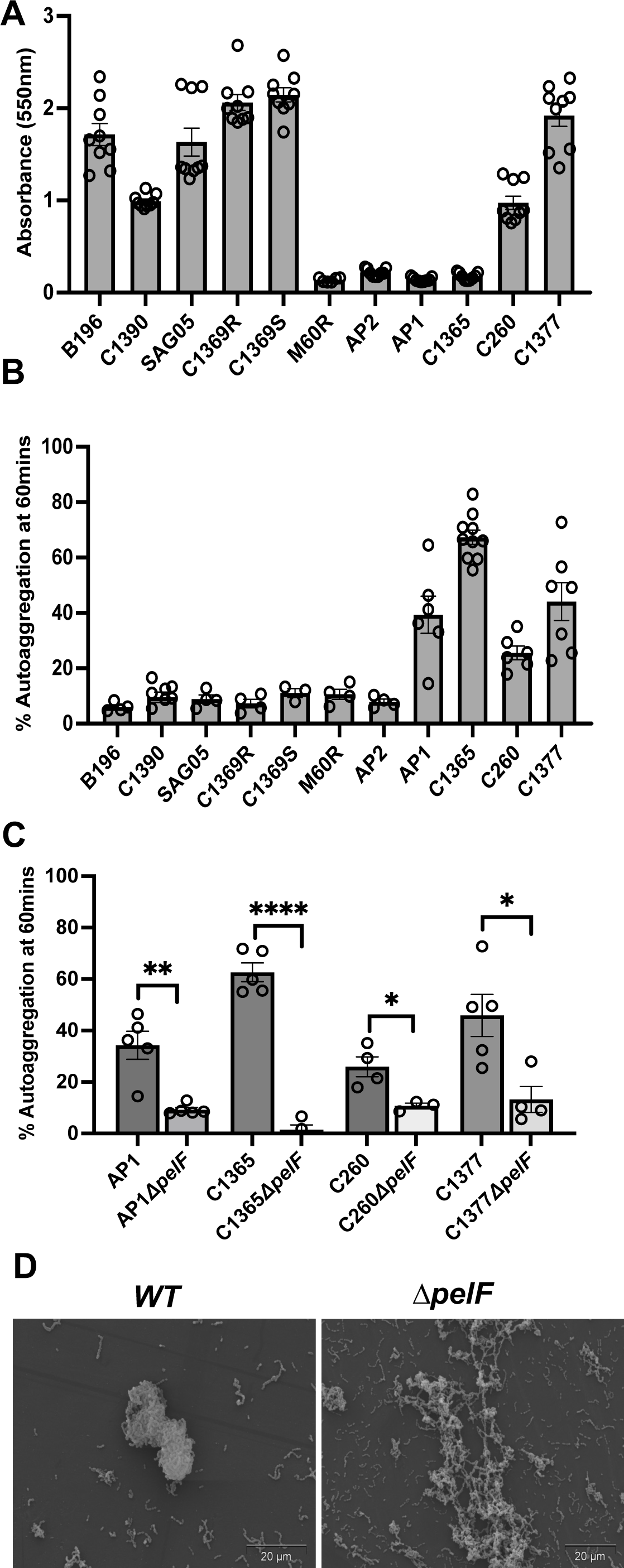
Pel biosynthesis contributes to auto-aggregation in hyper-aggregating strains of *S. intermedius*. **(A)** Surface adherence was assayed using crystal violet staining of biofilms formed in THY broth on poly-lysine coated polystyrene multi-well plates (Cellstar) after 24 hours, 37 °C, 5% CO_2_. **(B)** *S. intermedius* clinical strains were screened for auto-aggregation capacity at 60 min in an aggregation buffer. Four hyper-aggregating strains were identified. **(C)** Hyper-aggregation was abrogated by deletion of the glycosyl transferase *pelF* (*ΔpelF*) in the AP1, C1365, C260 and C1377, indicating that the process is dependent on Pel biosynthesis. Aggregation assays were performed at least 3 times. Aggregates formed by each wild type strain were compared to the corresponding *ΔpelF* mutant using an unpaired T test. * = p<0.05, **=p<0.01, ***=p<0.005, ****=p<0.001 **(D)** Scanning electron microscopy demonstrated that clumping of the hyper-aggregating strain *S. intermedius* C1365 is reduced dramatically in the *ΔpelF* strain.

### Auto aggregation in *S. intermedius* is dependent on the glycosyl transferase PelF

Bacterial aggregates have been linked to a variety of chronic infections and aggregation is known to be an important driver of adhesion and cohesion of biofilms in several bacterial species (8, 76). As surface adherence did not appear to be Pel dependent, we next screened the clinical isolates for their ability to auto-aggregate in suspension. We found four strains, AP1, C1365, C260 and C1377 displayed high levels of auto-aggregation over time in an aggregation buffer (Fig. 2B & Fig. S2A). Notably, deletion of the Pel glycosyl transferase *pelF* in each of these strains significantly reduced aggregation (Fig. 2C). Scanning electron microscopy (SEM) of the hyper aggregating C1365 strain corroborated the results observed in the solution assay as we found that the wildtype strain formed large cell aggregates with very few distinguishable chains of streptococci, while the cells in the C1365 *ΔpelF* strain were much more dispersed and chains of streptococci were clearly visible (Fig. 2D). Combined, these data suggest that auto-aggregation in *S. intermedius* requires the glycosyltransferase PelF.

### Auto aggregation in *S. intermedius* requires *pelD*

Two of the clinical *S. intermedius* strains, AP1 and AP2, were isolated from the pleural fluid of a patient with pneumonia. These strains displayed different phenotypes in our adherence and aggregation assays (Fig. 2A and 2B). While neither strain was surface adherent, AP1 aggregated, and this aggregation was dependent on the presence of *pelF.* Further characterization of these strains also revealed two different morphotypes. When grown on Congo-red agar plates AP1 colonies display a wrinkling around the circumference while AP2 had a smoother surface (Fig. S3A). To understand the origin of the phenotypic and morphologic differences, we sequenced the two strains. The sequencing revealed that the strains are isogenic except for three single nucleotide polymorphisms (SNPs). The first SNP is in the coding region of *glmU,* which encodes a *N*-acetylglucosamine-1-phosphate uridyltransferase that synthesizes UDP-*N*-acetylglucosamine (189). While the mutation results in a glycine to serine substitution at residue 304, examination of the AF2 model suggests that the UDP binding site and active site will not be affected, and that the function of this protein is unlikely to be disrupted. The second SNP is located 35 nucleotide base pairs upstream of a gene involved in copper homeostasis *cutC.* The third SNP is found in the *pelD* coding region. While the *pelD* gene is intact in AP1, the SNP in AP2 introduces a premature stop codon in the coding region of the third transmembrane helix of PelD. The premature stop would truncate the protein at residue 304 resulting in a protein that would lack the C-terminal GAF domain. Given our finding that deletion of *pelF* impacts aggregation in AP1 (Fig. 2B), we hypothesized that the *pelD* SNP in AP2 may result in the non-aggregation phenotype observed in this strain. *pelD* deletion mutants were therefore constructed in both AP1 and AP2 and their auto-aggregation phenotypes assessed. Deletion of *pelD* in AP1 resulted in a significant decrease in auto-aggregation relative to the wild-type parental strain while there was no detectable phenotypic change in the AP2 Δ*pelD* mutant (Fig. S3B). Taken together, these data suggest that PelD, like PelF, is required for the auto-aggregation observed in AP1, and that the SNP identified in AP2 reduces auto-aggregation to levels equivalent to that of a *pelD* deletion mutant.

### Pel production in *S. Intermedius* C1377 requires *pelDEA_DA_EF* but not *SIR_1591-1594*

Having demonstrated that both *pelD* and *pelF* are important for the aggregation phenotype, we next assessed the role of the rest of the genes in the *pel* operon in the hyper aggregating C1365 strain including the four additional non-canonical *pel* genes. Similar to the results obtained for the Δ*pelD* and Δ*pelF* mutants, deletion of each of *pelA_DA_, pelE* and *pelG* resulted in a loss of the aggregation phenotype (Fig 3A). Growth curves demonstrate that there were no apparent growth defects in any of these mutants (Fig. S4). Aggregation is still observed for each of the *SIR_1591, SIR_1592, SIR_1593* and *SIR_1594* deletion mutants, although there was a small but significant decrease in aggregation relative to wild-type observed for the Δ*SIR_1592* mutant (Fig. 3A). To determine whether this operon drives the production of a Pel-like polysaccharide, dot-blot analysis was performed using a monoclonal antibody (mAb) raised against a trimer of *N*-acetylgalactosamine (22, 77). This antibody is highly specific for the Pel polysaccharide. Our dot blot analysis clearly demonstrates that in the strains that aggregate, *i.e.* wild-type and each of the *SIR_1591, SIR_1592, SIR_1593* and *SIR_1594* mutants, the presence of Pel in cell pellets. As observed previously for *P. aeruginosa* and *B. cereus,* deletion of each of *pelDEA_DA_EF* results in the loss of Pel signal (Fig. 3B). While the Pel signal in the dot blot for Δ*SIR_1592* is suggestive of higher levels of Pel production in this mutant, quantification of this assay is difficult as OD optimization of the cultures for equal number of bacteria is difficult due to the aggregation phenotype. As analysis of *S. intermedius* C1365 aggregates using SEM did not reveal a visible biofilm matrix (Fig. 2C), confocal microscopy was performed using a fluorescently labelled version of the α-(GalNAc)_3_ mAb to verify that a Pel-like polymer was produced and its cellular location within the aggregates. Comparison of the C1365 wild-type and Δ*pelF* mutant clearly reveals localization of Pel to the large aggregates formed by the wild-type strain. No aggregates and no signal for Pel were observed in the Δ*pelF* mutant (Fig. 3C). Taken together these results suggest that each of the genes in the canonical *pelDEA_DA_EF pel* operon are required for the production of a Pel-like GalNAc-rich polymer that drives aggregation in *S. intermedius.* While deletion of any of the additional genes under the conditions tested did not abrogate aggregation or Pel production, there are subtle changes in the aggregation phenotype for SIR_1592 that suggest this protein may play a role in modulating the amount of Pel produced and/or its cellular location.

**Fig. 3.**
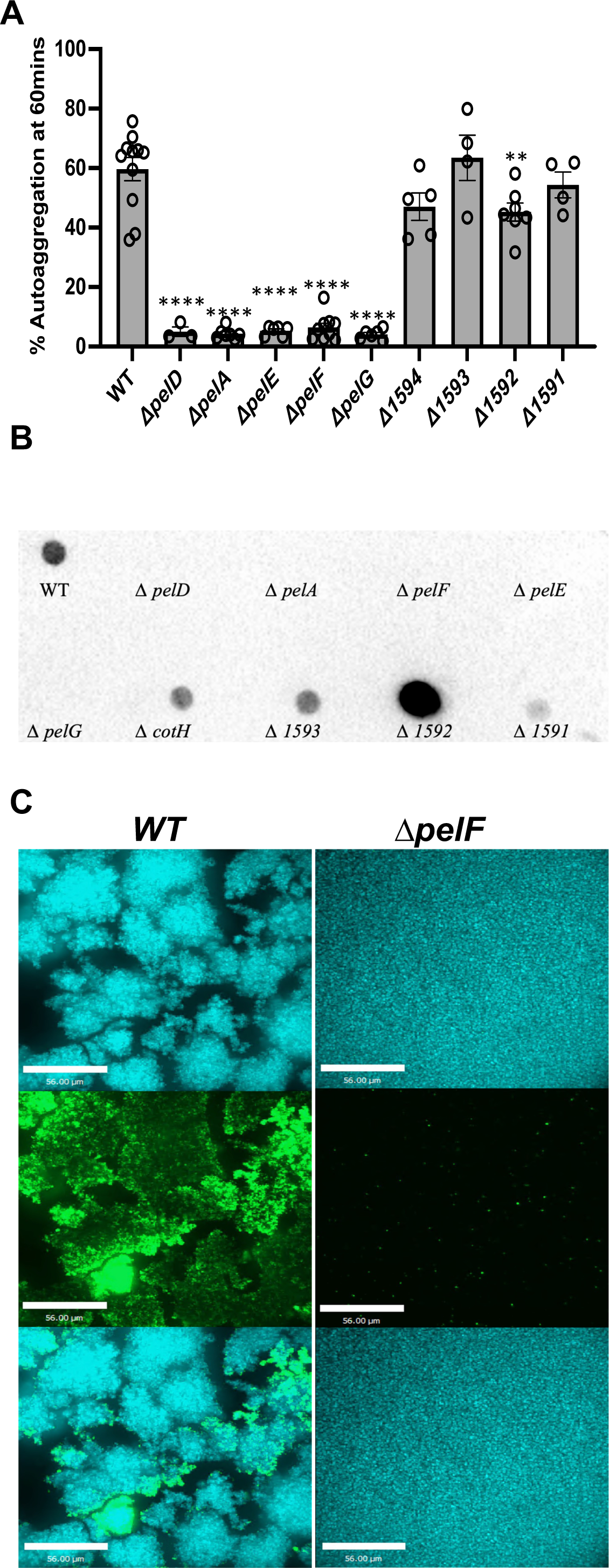
Contribution of *S. intermedius pel* genes to auto-aggregation. **(A)** Each of the genes in the *S. intermedius* C1365 *pel* gene cluster were deleted by allelic replacement and the deletion strains screened for auto-aggregation capacity. **(B)** *S. intermedius* C1365 wildtype and *pel*-deletion strains were assayed for presence of a Pel-like exopolysaccharide by dot blot. Cells were harvested and treated with proteinase K to remove protein, before probing for presence of a Pel-like polysaccharide using a monoclonal antibody raised against a BSA-(GalNAc)_3_ glycoconjugate. **(C)** Confocal images of *S. intermedius* C1365 wild type and Δ*pelF* deletion strain. Immunostaining was performed using a primary mouse anti-(GalNAc)_3_ antibody followed by secondary anti-mouse Alexafluor488 (Green) and DAPI to stain for bacterial cells (Blue). Images were collected using a Spinning Disk Confocal Microscope.

### Autoaggregation in *S intermedius* C1365 strain can be disrupted by the PelA hydrolase

To further characterize the Pel-like polysaccharide produced by *S. intermedius* C1365 aggregates, enzymatic disruption assays were performed. Cell aggregates were treated with enzymes that target different biofilm matrix components that have been shown to disrupt biofilm aggregates in other bacterial species (54, 56, 78–81). Using a panel of nine enzymes, including sugar-specific, protein-specific, and DNA-specific enzymes, we found that only the hydrolase domains of *P. aeruginosa* PelA, and *Aspergillus clavatus* Sph3 were able to disrupt *S. intermedius* C1365 aggregates (Fig. 4). The enzymes disrupted the aggregates with EC_50_’s in the nanomolar range. While both PelA and Sph3 have been characterized as α1,4-*N*-acetylgalactosaminidases, PelA can disrupt *P. aeruginosa* Pel-dependent biofilms, but Sph3 cannot (56). Differences in their active site architecture result in distinct enzyme-substrate interactions and their ability to hydrolyze deacetylated-rich regions within polymers. While neither enzyme can hydrolyze GalN oligosaccharides, PelA can hydrolyze partially deacetylated substrates better than Sph3. Of note, *Aspergillus fumigatus* Ega3, an endo acting α-1,4-galactosaminidase that can readily disrupt *P. aeruginosa* biofilms (78), did not disrupt the *S. intermedius* aggregates. These data provide further evidence that the hyper-aggregation observed in *S. intermedius* C1365 is dependent on an α1,4-*N*-acetylgalactosamine rich, Pel-like polysaccharide but suggest that there may be differences in the degree of de-*N-*acetylation of the polymer relative to *P. aeruginosa* or *B. cereus* (57, 78). The ability of Sph3 but not Ega3 to disrupt the aggregates suggests that *S. intermedius* Pel will predominantly be composed of GalNAc.

**Fig. 4.**
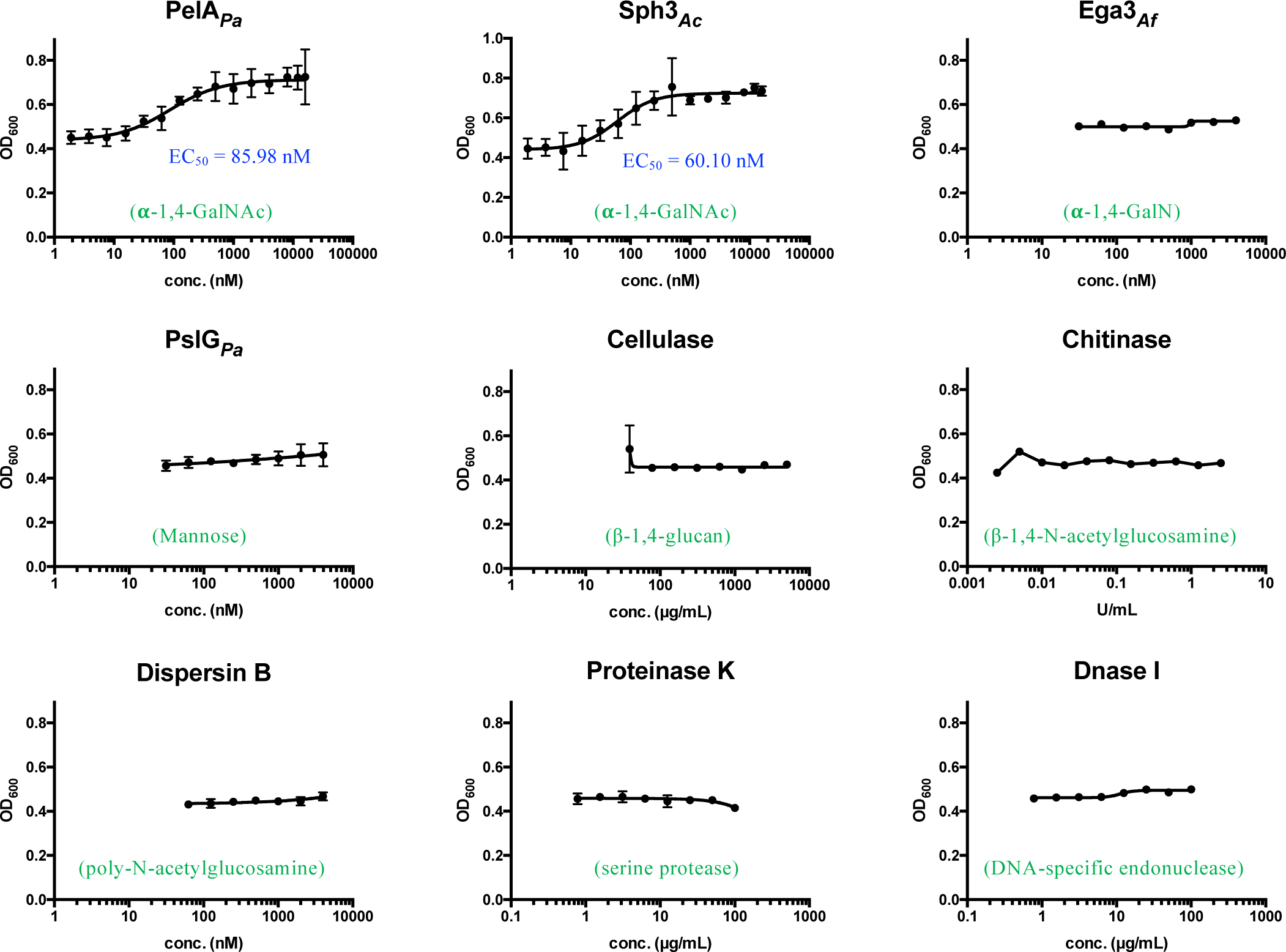
*S. intermedius* C1365 aggregation is driven by an exopolysaccharide chemically similar to *P. aeruginosa* Pel. Disruption assays were performed on *S. intermedius* C1365 aggregates, using a panel of enzymes targeting different matrix components. The graphs indicate dose-response curves to examine the disruption of aggregation by the exogenous treatment of each enzyme. Enzymes were added at a range of concentrations to bacterial aggregates and OD600 measured over time. EC_50_ values were calculated using nonlinear least-squares fitting to a dose-response model using Graph Pad Prism. Error bars indicate SEM.

### Pel dependent aggregation leads to longer lasting infections *in vivo*

Previous studies in *P. aeruginosa* have shown that the absence of cell free Pel enhances virulence in two infections models (19). Use of a Pel specific hydrolase in combination with Psl specific hydrolases and antibiotics reduces bacterial burden in both a lung model and a wound model of *P. aeruginosa* infection underlining the importance of the formation of biofilm aggregates and the role of exopolysaccharides in infection (57, 58, 82). To establish the importance of Pel and aggregate formation in virulence in *S. intermedius* we performed animal studies using a murine subcutaneous abscess model (83). Female BalB/C mice were infected with either the wildtype C1365 strain or the corresponding *ΔpelF* mutant sub dermally and the infection followed over six days. Abscess formation was examined at the site of injection and the size of the abscess measured daily. While we see visible abscesses in mice infected with the wild type strain, the mutant group had significantly smaller abscesses (Fig. 5 and S5). At the end of six days, the abscess size was measured, and the excised abscesses were weighed before homogenization and plating for bacterial burden. We found that one of the six mice in the mutant group had completely resolved the infection with no visible abscess after six days. The size and weight of the abscess were significantly lower in mice infected with the Pel deficient strain than in the wild type infected mice indicating that aggregates persisted for a longer time locally at the site of infection. We hypothesize that non aggregating bacteria disperse systemically and were rapidly cleared by the immune system. This hypothesis is also supported by the significantly reduced bacterial burden recovered from the abscesses of mice infected with the Δ*pelF* mutant (Fig. 5C).

**Fig. 5.**
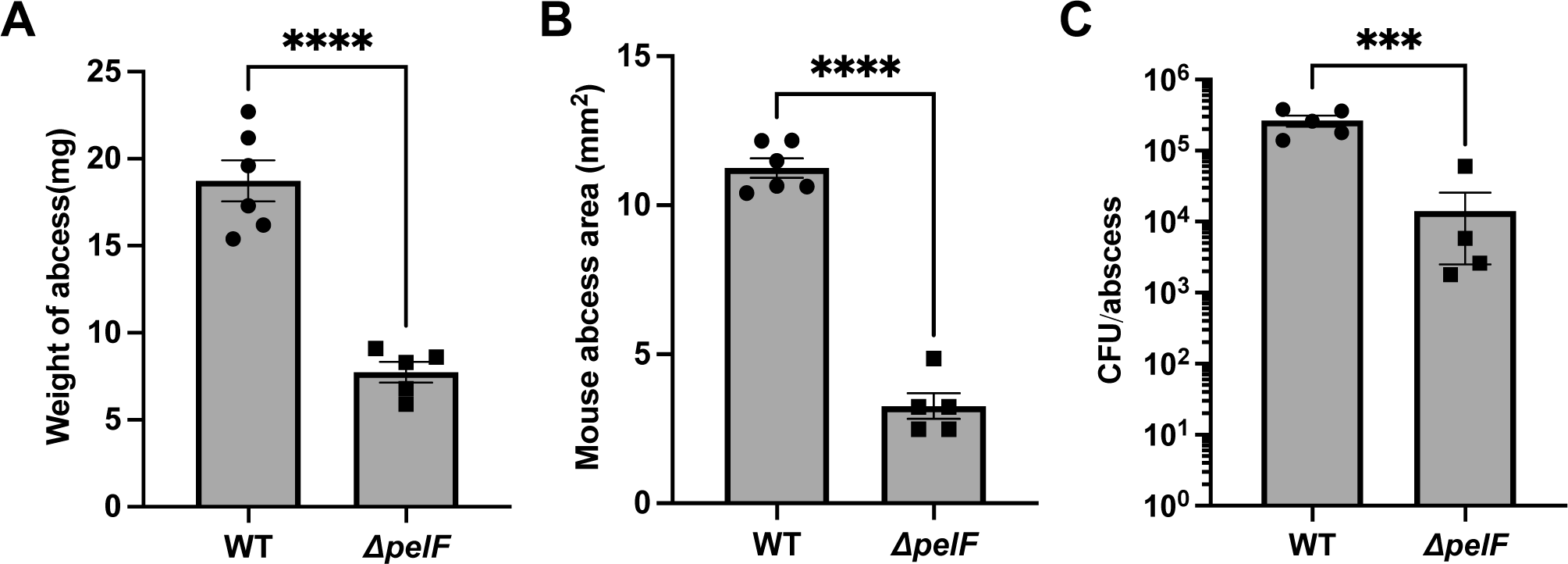
Lack of Pel allows for faster clearance of *S. intermedius* infections in a mouse abcess model. Female Balb/C mice were injected sub dermally with 1×10^8^ WT C1365 strain of *S. intermedius* and the corresponding *pelF* mutant. Infection was followed over 6 days for the development of abscesses. Abscess area was measured every day and the mice sacrificed at day 6 post infection. Weight of the abscess **(A)** and the abscess area **(B)** was measured post sacrifice and excision. Abscess homegenates were serially diluted and plated on THY agar. Plates were incubated for 24 h at 37 °C with 5% CO_2_ and CFU/abscess quantified. As observed in **(C)**, the CFU’s recovered from the mutant was significantly lower than from abscesses infected with WT Si. All statistical analyses were performed using unpaired T tests on Graph Pad Prism. ****=p<0.001, ***=p<0.005

### *S. intermedius* aggregation affects host immune response

Results from the murine abscess model indicate that Pel dependent aggregation is important for virulence of the pathogen and that these aggregates are not cleared by the immune system as efficiently as the non-aggregating *pelF* mutant. Induction of the immune response by bacteria is dependent on a myriad of factors. Predominantly, it is initiated when pattern recognition receptors sense conserved surface motifs on the bacterial surface or surface of the aggregates. This could include different components such as teichoic acids, lipopolysaccharides, surface proteins or the exopolysaccharides present on the surface of the aggregates. We hence hypothesized that the expression of extracellular polysaccharides such as Pel and the large aggregates formed by *S. intermedius* C1365 enable the bacteria to evade the innate immune response and alter cytokine responses. To test this, peripheral blood mononuclear cells (PBMCs) were isolated from four healthy donors and incubated with heat-killed *S. intermedius* cells *in vitro*. The cytokine profile of these PBMCs was then analysed by ELISA and compared to controls. We found that the average level of the IL-6, IL-5, IL-8, IL-10, GM-CSF, IL-1β, IFNγ and IL-12 cytokines were significantly increased in the hyper-aggregating C1365 strain compared with the non-aggregating C1390 strain (Fig. 6) and hypothesize that these differences could be because of the different biofilm matrix components in each of these strains. While the effects observed for the Δ*pelF* C1365 mutant were highly variable between donors, a significant reduction in the level of GM-CSF induced was observed in all donor PBMCs exposed to *S. intermedius* C1365 *ΔpelF* relative to the levels induced by exposure to the wildtype strain while levels of IL 12 were reduced in the wild type strain. These data support the hypothesis that hyper-aggregation affects how the *S. intermedius* cells interact with the host immune system. Further studies are needed to characterize the exact mechanisms by which the changes in the cytokine levels affect the immune response and thereby clearance of the bacteria from infection sites.

**Fig. 6.**
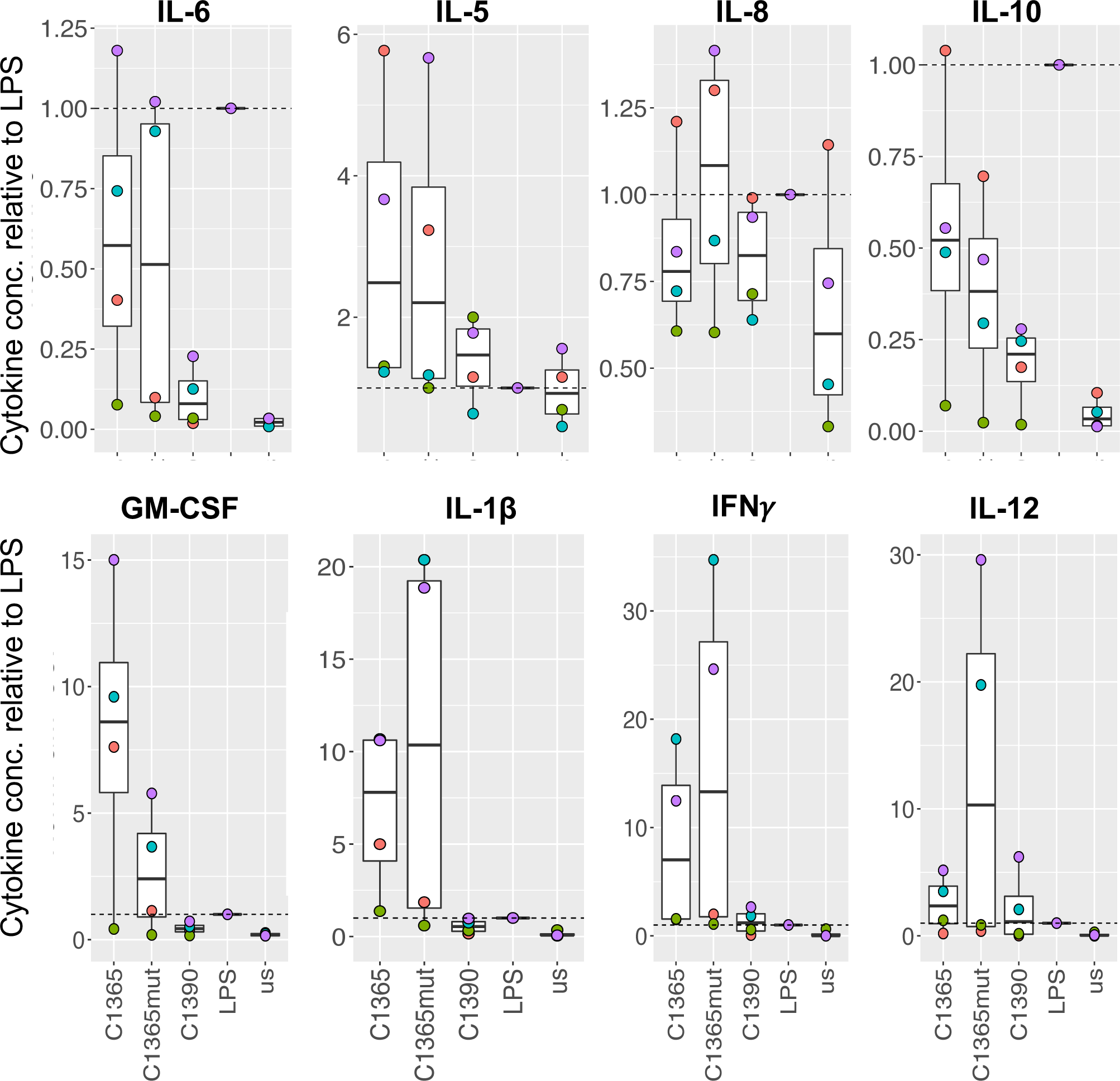
Hyper-aggregating strains of *S. intermedius* elicit a distinct GM-CSF and IL-12 response in human PBMCs. Heat-killed *S. intermedius* cells were incubated with primary human PBMCs isolated from 4 different donors, indicated by the coloured circles. Cytokine levels were normalized to LPS-induced controls to account for differences in the immunogenicity of the donors. us = unstimulated control.

## DISCUSSION

Herein we show that clinical strains of *S. intermedius* can auto-aggregate and that this aggregation is dependent on a Pel-like GalNAc-rich polysaccharide produced by *pelDEA_DA_FG* (Figs. 2-4). Interestingly, we found that surface adherence was not dependent on Pel (Fig. 2A). Animal studies using an abscess model show that Pel-dependent aggregation leads to longer lasting infections *in vivo* (Fig 5), while cytokine assays suggest that it can impact the interactions between the bacteria and the host immune system (Fig 6). Elevated levels of GM-CSF were detected from donor peripheral blood monocytes when exposed to the wild type C1365 Pel producing, aggregating strain. These GM-CSF levels were statistically different from the non-aggregating C1365 *ΔpelF* mutant or the non-aggregating C1390 *S. intermedius* strain. GM-CSF is an immune modulating cytokine that plays a critical role in maintaining the pulmonary immune system and is known to drive the immune functions of the alveolar macrophages (84, 85). GM-CSF negative mice have also been shown to clear Group B Streptococcal infections slower than wild type mice (85). While elevated levels of GM CSF would indicate an increase in the number of infiltrating macrophages, it has been shown previously that the beneficial impacts of bacterial aggregation are more pronounced for larger aggregates as larger aggregates are more resistant to phagocytosis (86). Thus, an elevated immune response to aggregates could lead to more harm than good by increasing inflammation levels. The cytokine, IL-12, also showed significant differences in PBMC’s from two donors. The IL-12 response in aggregating strains was muted compared to the response in the *ΔpelF* strains. IL-12 levels are critical in early infection control, generation and maintenance of an adaptive immune response and phagocytosis (87). These results are possibly indicative of the ability of the host immune response to more effectively clear Pel deficient non aggregating strains. While we did not observe significant changes in other cytokine levels, this study only assessed the role of PBMC’s. Futures studies will need to explore whether other aspects of the immune response control the host’s response to infections.

While the current study focussed on the Pel dependent aggregating strains of *S. intermedius,* our analyses of eleven different clinical isolates enables us to define four different phenotypes based on their ability to form surface adherent and/or aggregating biofilms. The phenotypes observed were: (i) aggregating and surface adherent (C1377, C260); (ii) aggregating and non-surface adherent (C1365, AP1); (iii) non-aggregating and surface adherent (B196, SAG05, C1369S, C1369R, C1390,); and (iv) non-aggregating and non-surface adherent (M60R, AP2). Even though our data clearly support a role for Pel in aggregation, it also suggests a limited role for Pel in surface adherence. These data contrast with observations for *P. aeruginosa* and *B. cereus* where Pel has been shown to play a role in adherence in these species (16, 21). The limited role for Pel in *S. intermedius* adherence could be related to structure and therefore charge of the mature Pel polymer. Our aggregate disruption assays revealed that the α-1,4-*N-* acetylgalactosaminidases PelA and Sph3, but not the α-1,4-galactosaminidase Ega3, disrupt *S. intermedius* aggregates. These data suggest that cell-associated *S. intermedius* Pel may be composed predominantly of GalNAc, which would lead to a neutral rather than cationic polymer. Differences in the charge of the polymer will have a profound effect on its function. The structure of *S. intermedius* Pel and what other factor(s) in the biofilm matrix of these clinical strains contribute to the different phenotypes, especially adherence, remains to be determined.

Previous analyses of the *pel* operon in different bacterial species (15–17) identified *pelD* as a key difference between the *S. intermedius,* and *P. aeruginosa* and *B. cereus pel* operons. PelD has been classified into 4 classes based on the presence or absence of the GGDEF, SDR or degenerate SDR domains. *S. intermedius* PelD falls into Class IV, as it contains a degenerate SDR domain but lacks the GGDEF domain and hence does not have the ability to bind c-di-GMP (17). Bioinformatic analysis of available SMG genome sequences also indicate that these species do not encode homologues to any known diguanylate cyclase enzymes. This suggests that unlike *P. aeruginosa* Pel biosynthesis, which is regulated at the transcriptional and protein levels *via* binding of c-di-GMP to FleQ and PelD, respectively (33, 34, 88), transcription of the *pel* operon and the activation of the *S. intermedius* Pel biosynthetic complex must be regulated by different mechanisms. Our gene cluster analysis reveals *pel* operons across multiple SMG species (Table S2) providing strong evidence to suggest that this operon plays an important role in the biology of these species, not just *S. intermedius,* and that *pelD* is a functional part of the system. The importance of PelD in auto-aggregation is clearly demonstrated by the AP1 and AP2 clinical isolates (Fig S3). How Pel biosynthesis is regulated in the absence of c-di-GMP is yet to be determined but c-di-AMP and ppGpp are potential candidates as in other Gram-positive species, these molecules have been shown to play a role in regulating the response to nutrient limitation and biofilm formation (89–93). Given the multiple levels of Pel regulation in Gram-negative bacteria, it will be interesting to study how in the absence of c-di-GMP, other nucleotides regulate the *pel* operon and its expression in *Streptococci*.

The other notable difference when comparing the *S. intermedius pel* operon to the operons found in *P. aeruginosa* or *B. cereus* is the presence of the four additional genes. Our results show that deletion of each of these additional genes under the conditions tested did not affect the ability to make Pel or aggregate. This was surprising as *SIR_1591* is predicted to be a glycoside hydrolase and deletion of the hydrolase adjacent to the *pel* operon in *B. cereus* led to an increase in biofilm formation (16, 19). We had anticipated that we would see a similar phenotype but deletion of *SIR_1591* in *S. intermedius* did not increase either aggregation or surface adherence (Fig. 3A). Similarly, systematic deletion of the other additional genes did not cause significant changes in either aggregation or Pel production. However, our analyses found that *SIR_1592-1594* always occur together within a *pel* operon (Table S2) suggesting that this combination of genes are important. Interestingly, *SIR_1592* and *SIR_1593* are both connected to the poly-phosphate synthesis pathway and functional homologues of these genes seem to play major roles in host pathogen interactions and stress responses in bacteria (71, 74, 94). Given that most of our experiments were conducted in rich media in a host independent system, it is possible that the functional relevance of these additional genes and how they relate to Pel production, and Pel-dependent phenotypes have not been fully explored. Future studies will need to study the role of these genes in the context of the host using, for example, epithelial or macrophage cell lines.

This study establishes a role for Pel in aggregation and in infection in SMG organisms, specifically *S. intermedius*. We show that Pel leads to persistent infections *in vivo* and can modulate host immune responses. Our analysis of the operon highlights the differences between Streptococcal spp and other Gram-positive and -negative species such as *B. cereus* and *P. aeruginosa.* Future studies will focus on determining the c-di-GMP independent mechanism of regulation of Pel in these bacteria, and the precise role of the four additional genes.

## Data Availability

All data generated or analyzed during this current study are included in this published article and its supporting information.

## Acknowledgements

We thank the staff of the Nanoscale Biomedical Imaging Facility at The Hospital of Sick Children for assistance with preparation of the SEM samples.

## Contributor roles

**Conceptualization:** DR, SAT, PLH

**Investigation:** DR, SAT, KC, GBW, DM, SM, RP,

**Resources:** FM, DCS, MJ, SS, TLL

**Project administration:** PLH

**Supervision:** TLL, MS, DCS, DJW, PLH

**Writing – original draft:** SB, DR, RP, PLH.

**Writing – review & editing:** All authors reviewed and approved the final manuscript.

