## Supplementary material for "Autoaggregation in *Streptococcus intermedius* is driven by the Pel polysaccharide": Table S1 and SI Figures

### Table S1: Primers used in this study

| Primers | Sequence* |
|---|---|
| **Allelic exchange vectors** | |
| KanR F | GGTCCGATAAACCCAGCGA |
| KanR R | CGATACAAATTCCTCGTAGGC |
| PelD up F | GGG**TCTAGA**TGGCTATTTTTGGAGACGGG |
| PelD up R | TCGCTGGGTTTATCGGACCCCCTGTATTTCCAAATAGTAGAG |
| PelD down F | GCCTACGAGGAATTTGTATCGACTCCTAATGGCATCAAATATGG |
| PelD downR | GGG**GGATCC**ACAAAACTGCAGCTAATAAAAGAAA |
| PelE up F | GTAG**GGATCC**CATCTGGAATTCAATCGTTTAAAAGC |
| PelE up R | GTTCGCTGGGTTTATCGGACCTGCCAAATCTAAAATCACACAGAATAG |
| PelE down F | CCTACGAGGAATTTGTATCGTCAAAAGAGAACCGAGAATATTTAGCT |
| PelE downR | GTCA**CTCGAG**CAGATAACTTCAATAAATCTGGCC |
| PelA up F | GTAG**GGATCC**ATCATCAAGCATTTGTCAACCTC |
| PelA up R | GTTCGCTGGGTTTATCGGACCCTCCCTTGAATAAGCCAATACC |
| PelA down F | CCTACGAGGAATTTGTATCGGCTACTTCTGCAAAAATAGAAATTG |
| PelA downR | GTCA**CTCGAG**GAACATTGGGCATCTGGGAG |
| PelF up F | GGGGAGCTCTATTTAGACGATTTTCCTTCCCCT |
| PelF up R | GTTCGCTGGGTTTATCGGACCACCCTCTAAAACCAAACAGATTCT |
| PelF down F | CCTACGAGGAATTTGTATCGAGACAATTATATAAGGAGTATGTAAGA |
| PelF down R | AAGGGGCATGCGTAAAACTGTCAACAGAACAATTC |
| PelG up F | GTAG**GGATCC**CATACTATCCGATCTATGCTTTTTCC |
| PelG up R | GTTCGCTGGGTTTATCGGACCTCGCAGTTCGAATCCTATCC |
| PelG downF | CCTACGAGGAATTTGTATCGGTGACTATTTTGAAAGCGAGGTC |
| PelG downR | GTCA**CTCGAG**CCATCAATATTTGAAAGGCCAACC |
| 1592 up F | GTAG**GGATCC**GGTCTAACAAAGACAAGTGTTACG |
| 1592 up R | GTTCGCTGGGTTTATCGGACCTGTAAAATTTTCCGATCTCAAAACCG |
| 1592 downF | CCTACGAGGAATTTGTATCGCAGTATAATGGAGAGTATCACGG |
| 1592 downR | GTCA**CTCGAG**GGACTTCAGCAATTTTAGGC |
| 1593 up F | GTAG**CATATG**TTCGAAAGGAAAATGATCCTAGG |
| 1593 up R | GTTCGCTGGGTTTATCGGACCCTGTTCTTTTTTATTTGCTTTCATAGAC |
| 1593 downF | CCTACGAGGAATTTGTATCGCATGATTTTGAATATTGGTATGATACAGATC |
| 1593 downR | GTCA**CTCGAG**CGTTTTCCAATCATAGACTCC |
| 1594 up F | CTA**GTCGAC**GAAAGATTATCGTAGTATTCTAAAAACCTTTGCCAT |
| 1594 up R | GTTCGCTGGGTTTATCGGACCTAAAAATTTGATCCACTTATTATCCATCTTTTGACC |
| 1594 downF | GCGCCTACGAGGAATTTGTATCGATAAAGGTTGATGTCTATGAAAGCAAATAAAAAAGAAC |
| 1594 downR | CTA**GGTACCG**ATTCGAACTAATGGCCAAAGTAACCATAGC |
| **Sequencing Primers** | |
| PelD F | CCA AGC TTG CAT GCC TGC AG |
| PelD R | CAG TCA CGA CGT TGT AAA ACG AC |
| PelE F | GGCACCGTTATGGCTGGTTT |
| PelE R | GCAGTTACATGACCTGATGTATCATC |
| PelA F | ATGATTGGTGGAGCAACAGC |
| PelA R | TGCTAAAGCATAACATTCTTCTGC |
| PelF F | TATGAAAGAGCAGTTCGCGGTAT |
| PelF R | ACCCCATAGCCTAAACAAAGTG |
| PelG F | TTTTCGACCTCCTTCAAGAAGG |
| PelG R | TATACCAACGTTTCCCATGTAGG |
| 1594 F | CTTGATCTTGTTTAGTGAACTATG |
| 1594 R | ATCCTTAACTGCTGTCCGGT |
| 1593 F | GCTCTTGAAGCACATGGACTAG |
| 1593 R | CCTTCTTTGCGCCAGCGATT |
| 1592 F | TCATGTGGAGTTAGCTAAAGATCC |
| 1592 R | GTAACTGACAGTGTAAGCTTAGTGG |
| 1591 F | AGTCCTCATGCCTTGTGTCGT |
| 1591 R | GCAAAACCAGCCAAAAGAGCACTC |
| T7 | TAATACGACTCACTATAGGG |
| T7 ter | GCTAGTTATTGCTCAGCGG |

| **Plasmid Vectors** | |
|---|---|
| pET28a | IPTG inducible protein expression vector- Kanamycin resistant |
| pCC1-4k | Plasmid vector used by Biobasic for inserting 1591 mutant allele- Chloramphenicol resistant |

1

* Restriction sites are indicated in bold and overlap with Kan cassette in the allelic exchange vectors are underlined

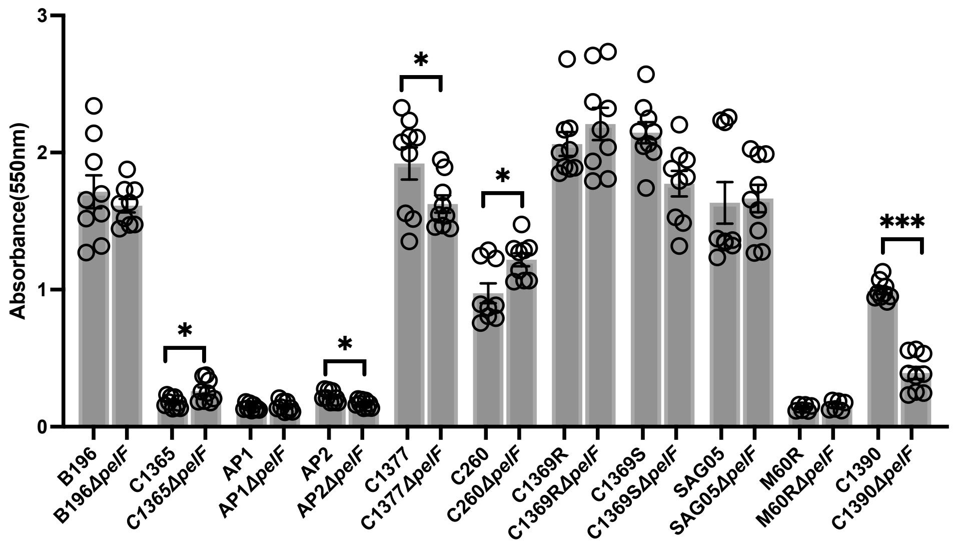


**Supplementary Figure 1**. **Pel does not significantly affect surface adherence in *S. intermedius.*** Eleven *S. intermedius* clinical strains and the corresponding Δ*pelF* mutants were screened for their ability to adhere to 96 well tissue culture plates using a crystal violet adherence assay.

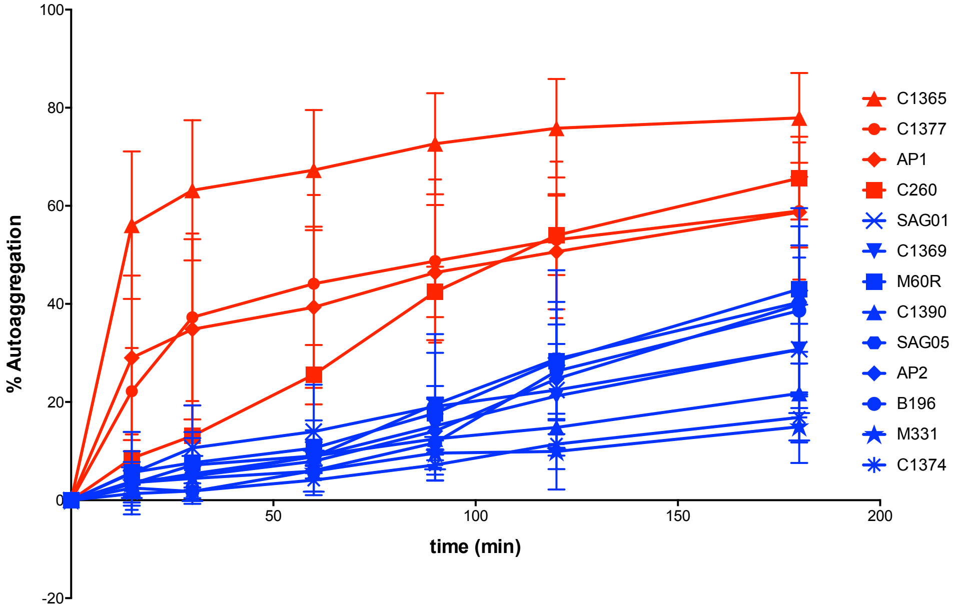


**Supplementary Figure 2. Pel biosynthesis contributes to auto-aggregation in hyper-aggregating strains of *S. intermedius***. *S. intermedius* strains were screened over time for auto-aggregation capacity. Four hyper-aggregating strains were identified (red).

**A**

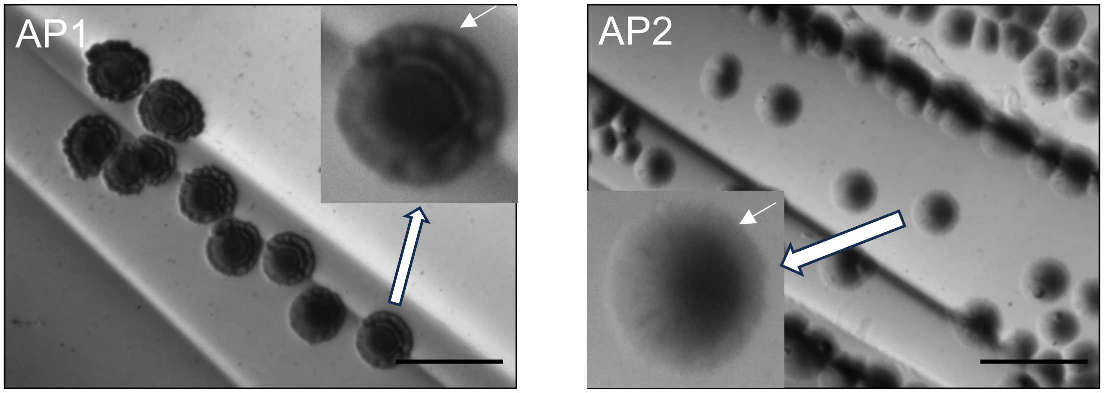


**B**

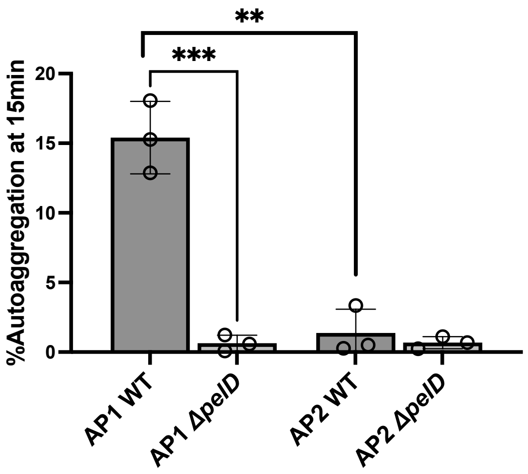


**Supplementary figure 3: Truncation of *pelD* gene in AP2 results in a loss of the aggregation phenotype.** (**A**) Colony morphology of clinical isolates, AP1 and AP2, observed under a stereoscopic microscope. The strains were streaked onto THY-congo red agar and incubated at 37 ºC with 5% $CO_2$ for 3 days. AP1 displays a wrinkly phenotype while AP2 has a smoother surface. The scale bar is equivalent to 5 mm. Inset shows a zoomed in colony in each panel with the small white arrows indicating the wrinkly vs smooth edges in AP1 and AP2 respectively. (**B**) An aggregation assay comparing phenotypes of the clinical isolates AP1 and AP2 at 15 min. AP1 wild- type (WT) auto-aggregates to a greater extent than WT AP2. Deletion of *pelD* (Δ*pelD*) in AP1 results in a decrease of auto-aggregation compared to the AP1 WT strain at 15 min. Low auto-aggregation is observed in both the AP2 WT and AP2 Δ*pelD* strains. Error bars represent standard deviations from three independent samples. Statistical analyses were performed on 3 biological replicates using unpaired T test to compare each wild type strain to its derived mutant. AP1 and AP2 were compared separately using a T test. ** $=p \leq 0.01$, ***$= p<0.005$

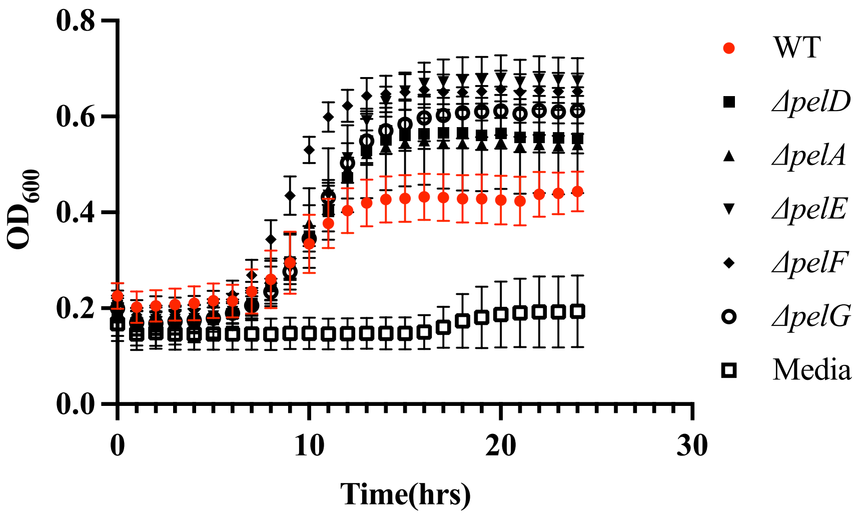


**Supplementary figure 4. Growth curves for wildtype and *pel* gene cluster mutant strains of *S. intermedius* C1365.** *S. intermedius* wildtype strain C1365 (WT) and the corresponding mutants of the genes in the putative *pel* operon were grown in THY broth overnight at 37 ºC and 5% $CO_2$ in a 96 well plate. The plate was incubated in a SpectraMax i3X Multi-Mode Assay Microplate Reader. Optical density (600nm) was measured every hour over 24 h. There were no significant differences observed in the growth between the WT and the mutants.

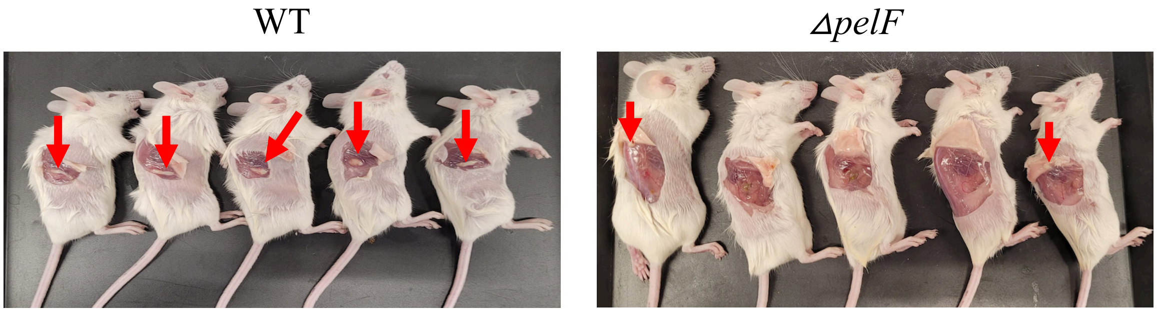


**Supplementary figure 5. Murine subcutaneous abscess model for pathogenicity of *S. intermedius* aggregates**. Female Balb/C mice were infected with wildtype C1365 (WT) or with the corresponding PelF mutant(*ΔpelF)*. Mice were sacrificed 6 days post infection and the abscesses measured for size, weight and bacterial burden. Red arrows show presence of abscess in the infected mice after 6 days.
